# Population structure and clonal prevalence of scleractinian corals (*Montipora capitata* and *Porites compressa*) in Kaneohe Bay, Oahu

**DOI:** 10.1101/2019.12.11.860585

**Authors:** NS Locatelli, JA Drew

## Abstract

As the effects of anthropogenic climate change grow, mass coral bleaching events are expected to increase in severity and extent. Much research has focused on the environmental stressors themselves, symbiotic community compositions, and transcriptomics of the coral host. Globally, fine-scale population structure of corals is understudied. This study reports patterns of population structure and clonal prevalence found in *Montipora capitata* and *Porites compressa* in Kaneohe Bay, Oahu. Generated using ddRAD methods, genetic data reveals different patterns in each taxa despite them being exposed to the same environmental conditions. STRUCTURE and site-level pairwise F_ST_ analyses suggest population structure in *M. capitata* resembling isolation by distance. Mantel tests show strong, significant F_ST_ correlations in *M. capitata* in relation to geographic distance, water residence time, and salinity and temperature variability (range) at different time scales. STRUCTURE did not reveal strong population structure in *P. compressa.* F_ST_ correlation was found in *P. compressa* in relation to yearly average sea surface height. We also report high prevalence of clonal colonies in *P. compressa* in outer bay sites exposed to storms and high energy swells. Amongst only outer bay sites, 7 out of 23 sequenced individuals were clones of other colonies. Amongst all 47 sequenced *P. compressa* individuals, 8 were clones. Only one clone was detected in *M. capitata*. Moving forward, it is crucial to consider these preexisting patterns relating to genetic diversity when planning and executing conservation and restoration initiatives. Recognizing that there are differences in population structure and diversity between coral taxa, even on such small-scales, is important as it suggests that small-scale reefs must be managed by species rather than by geography.

## Introduction

Rapid climate change due to anthropogenic carbon emissions is one of the greatest threats to global marine biodiversity (Cheung et al. 2009). Within the past few decades, coral bleaching events have increased in occurrence and severity to the point where they are becoming commonplace (Hughes et al. 2003). Despite bleaching being a widely-known impact of climate change, the pathways by which it occurs remain poorly understood.

A large proportion of research has focused on the role of zooxanthellae, dinoflagellate algae of the genus *Symbiodinium* that form symbiotic relationships with coral, in mediating the bleaching response. In a zooxanthellae driven response, thermal bleaching is caused by or begins when photosystems within the symbiont cells become damaged by heat and sunlight and cells are subsequently ejected by the coral host (Jones et al. 1998, Warner et al. 1999). In addition to symbiont-related mechanisms of coral bleaching, bleaching can be a physiological response of the coral, in which case genetic variation among coral could affect their response. Some evidence exists for this mechanism. When experimentally exposed to warm water, populations of *Porites astreoides* from different temperature conditions (no more than 10km apart) showed different bleaching responses despite harboring the same *Symbiodinium* communities. These responses were associated with differences in gene expression and significant genetic divergence correlated with *in situ* temperature conditions (Kenkel et al. 2013, Kenkel and Matz 2016). A third mechanism for coral bleaching is the probiotic hypothesis (Reshef et al. 2006). This mechanism has highlighted the importance of microbial communities in coral mucus and tissues that change in response to abiotic conditions such as temperature (Bourne et al. 2008, Li et al. 2015). Studies have shown that increasing water temperature is associated with a shift in bacterial community compositions and virulence patterns and that following temperature stress, bacterial communities slowly return to their original state (Bourne et al. 2008, Rosenberg et al. 2009).

These mechanisms are usually studied separately and do not consider the effect of population dynamics of the coral host. This oversight may be partly due to the difficulty of studying population genetics in many coral genera until the recent application of restriction-site associated methods, primarily in Caribbean corals (Drury et al. 2016, 2017, Devlin-Durante and Baums 2017, Forsman et al. 2017). Although microbial communities are essential to the long-term survival of corals, studying these communities without considering the genetic structuring of the coral host leads to an incomplete understanding of the drivers of bleaching events. This study seeks to understand population genetic structuring patterns of two Pacific reef-building corals, *Montipora capitata* and *Porites compressa*, in Kaneohe Bay, Oahu.

*M. capitata* and *P. compressa* were chosen as the focal species due to their wide ranges and their importance as major reef-building organisms in shallow waters of the Main Hawaiian Islands. *Montipora* are generalists in their *Symbiodinium* community composition but are generally more sensitive to environmental conditions than *Porites*, which are largely inflexible to shifting symbiont composition (Putnam et al. 2012). Growth rates differ between the taxa, with *Montipora* having high growth rates and *Porites* a comparatively low rate (Gladfelter et al. 1978, Huston 1985). This suggests that there may be an inherent fitness tradeoff associated with symbiont switching ability. *Montipora* switch symbionts to optimize for fast growth at the expense of environmental sensitivity while *Porites* exhibit high symbiont fidelity that confers environmental resilience but slower growth. Because of this inherent difference, it is imperative for the field to better understand if these corals, with fundamentally different life history strategies, differ in their genetic structure.

Kaneohe Bay is a well-studied marine system that is uniquely positioned to explore these questions. The Hawaii Institute of Marine Biology (HIMB) sits upon Coconut Island in the southern, sheltered portion of the bay and is the gateway for much of the research that comes out of the bay. As a result of this, episodes of extreme stress, like heatwaves and freshwater kills, are well-documented and the patterns of bleaching in 1996 and 2014 documented by researchers at HIMB provide some context for this present study (Jokiel and Brown 2004, Bahr et al. 2015a, 2017). In addition to its recent temperature-related stressors, the bay has a long history of human utilization that began with Polynesian settlement and has more recently been subject to invasive species introduction, agricultural runoff, sewage discharge, and extensive dredge and fill operations (Bahr et al. 2015b). In an otherwise well-studied system, the bay is understudied in regards to the population genetics of their hallmark organisms: corals. This study sought to fill this gap in knowledge by utilizing ddRAD (Peterson et al. 2012) to understand the population structure and genetic diversity of corals within Kaneohe Bay (KB) and determine if any patterns differ between the sampled taxa.

## Methods

### Sample Acquisition and Preservation

Colonies of *M. capitata* and *P. compressa* were collected between August 20^th^ and August 30^th^ of 2018 under authorization from the Hawai’i Department of Land and Natural Resources Special Activity Permit (SAP) No. 2019-67. A total of 48 individuals from each species were collected from among eight sites (six individuals per site per species). Sites were evenly spaced to capture the diversity of abiotic conditions found throughout KB (**Fig. 1**, sample inventory and GPS coordinates in **Supplemental Table 1**). Samples were obtained from colonies <1m^3^ in size growing outside of prohibited areas and monitored reefs as outlined in the SAP. Sites were 1-2m in depth with the exception of site 6 which was ∼5m in depth. Colonies were sampled >20m apart to minimize the possibility of fragmentary clones. Fragments ∼2cm in length were taken from each individual and immediately preserved in 100% ethanol in 1.5ml microcentrifuge tubes. Samples were shipped to the continental US, transferred to sterile 5ml microcentrifuge tubes, and topped off with additional 100% ethanol to increase the overall ethanol concentration for long-term storage.

**Figure 1:**
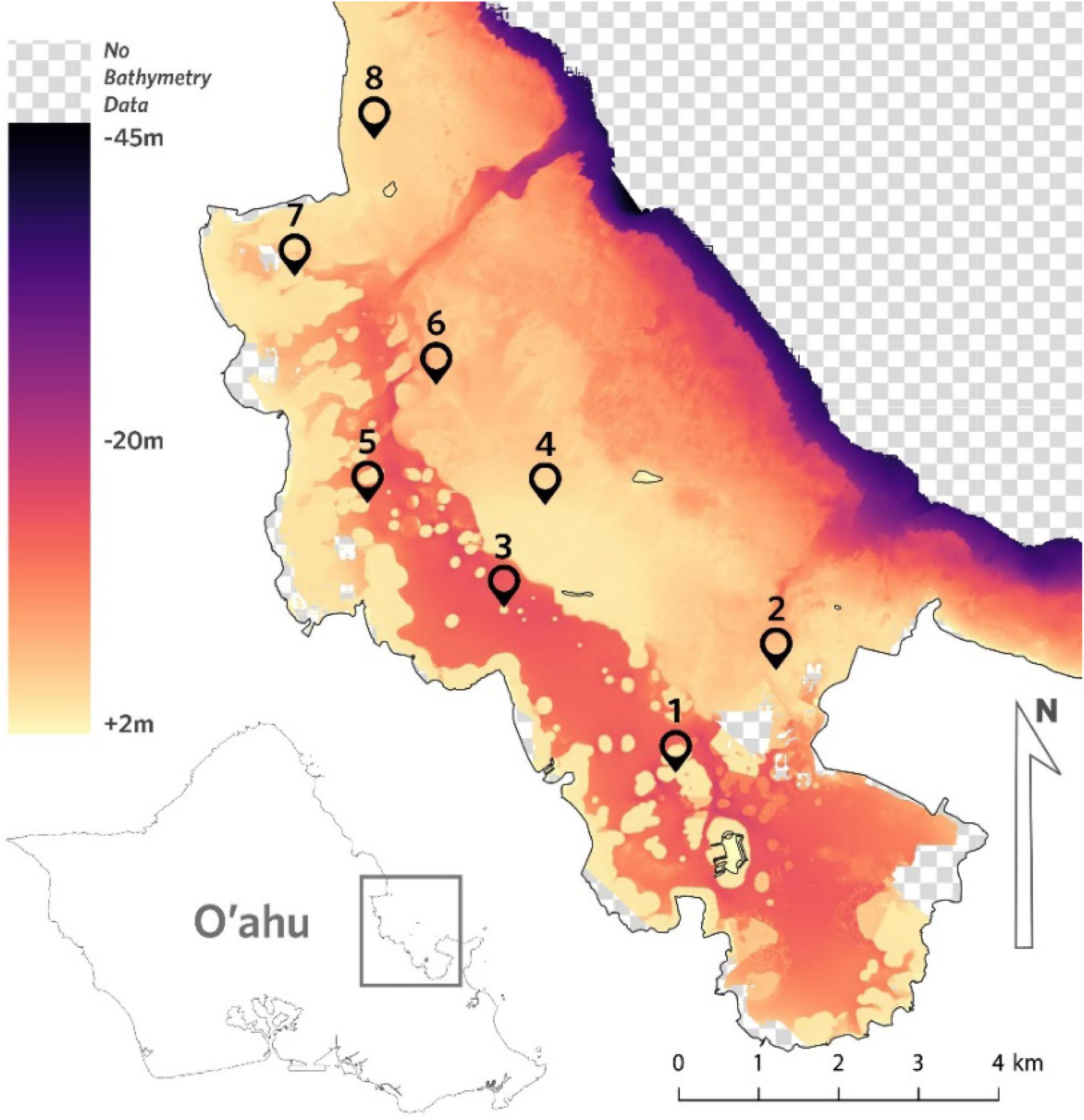
Sampling sites within Kaneohe Bay, Oahu. Sites ranged from 1-5m in depth and six individuals per site per species were collected for *Montipora capitata* and *Porites compressa*.

### DNA Extraction, Library Preparation, and Sequencing

Ethanol-preserved samples were placed on sterile mixed cellulose ester (MCE) membrane filter to absorb and evaporate excess ethanol. Tissue was obtained by removing the outermost ∼1mm of material from a surface area of approximately 1cm^2^ using a sterile scalpel. Removed tissue and skeletal material was pulverized in the folded MCE membrane using the side of the scalpel blade. Pulverized tissue was allowed to dry completely and transferred to a 1.5ml microcentrifuge tube. For *Montipora* samples, extractions were performed using E.Z.N.A. Tissue DNA Kits (Omega Bio-Tek) with unmodified protocols. For *Porites* samples, extractions were performed with unaltered protocols with the exception of centrifugation steps. Excessive mucopolysaccharides severely clogged spin columns and required additional time and velocity (20800RCF) to push the fluid through silica columns. Extracts were quantified using an AccuGreen™ Broad Range dsDNA Quantitation Kit (Biotium) with a Qubit 3.0 fluorometer (Invitrogen). Yields of *Montipora* extractions were all >65ng/microliter while yields for *Porites* samples ranged from 3.96ng/µl to 106ng/µl. Volumes of eluted DNA ranged from 100-250µl. Salt-ethanol precipitations using 3M sodium acetate (pH ∼7.0) were performed on low concentration samples such that all met the library preparation and sequencing provider requirements of >25ng/µl concentration and >20µl volume. No laboratory methods were used to minimize symbiont contamination from Symbiodinaceae symbionts. Contamination in *Montipora* samples were anticipated to be low as fragments were taken from apical growing tips which hold low concentrations of symbionts (Oliver 1984). Based on color of sample and solution, most symbiont cells in *Porites* samples were thought to have been present in solution and low contamination was expected.

Extracted DNA was sent to the University of Minnesota Genomics Center for the Sequence-based Genotyping (SBG) service for library preparation and sequencing. The 96 samples underwent quality control and re-quantification to verify sufficient sample volume and mass. The library preparation method utilized was ddRAD (Peterson et al. 2012) using TaqI and BtgI as restriction enzymes. At the time of enzyme selection, a *Montipora capitata* genome was not yet available. Thus, enzymes were chosen based on an expected genome length of 420-552Mb as inferred from published *Acropora digitifera* and *Porites lutea* genomes. Fragments were subsequently size selected for the range of 300-744bp with an insert size of 156-600bp and then amplified. Prepared fragments were sequenced using half a lane of NextSeq 500 in high output configuration with single-end chemistry (1×150bp).

### Data Processing and Bioinformatics

Demultiplexed data was received from the University of Minnesota Genomics Center and preliminary quality control analysis was performed using FastQC (bioinformatics.babraham.ac.uk/projects/fastqc/). Small amounts of Nextera Transposase adapter sequences were detected and the first 15bp of each sequence were biased in their content. Trim Galore (bioinformatics.babraham.ac.uk/projects/trim_galore/) was used to remove remaining adapters as well as the first 15bp of each read. Because no outgroup was sequenced, *M. spumosa* sequence data (Consortium 2015) was in-silico digested using FRAGMATIC (Chafin et al. 2018), duplicated 10x to increase “read” depth, converted from fasta to fastq using dummy quality scores, and included in the assembly. No in-silico data was needed for *P. compressa* as ipyrad allows for the inclusion of reference genotypes (*P. lutea* reference genome) as a sample in output files. Following quality checks, data for each species was assembled using the ipyrad 0.9.4 pipeline (Eaton 2014, ipyrad.readthedocs.io). Assembly of *M. capitata* reads was performed using the newly published *M. capitata* nuclear genome assembly (Shumaker et al. 2019) and assembly of *P. compressa* reads was performed using the *Porites lutea* genome assembly (ReFuGe 2020 Consortium, Liew et al. 2016). The datatype selected was single-end ddRAD, assembly type was “reference”, and all formats were output. All other parameters were left as default.

PHYLIP alignments of variant sites generated from the ipyrad assembly were input into RAxML-NG (Kozlov et al. 2019) to generate a maximum likelihood tree with bootstrap support for *M. capitata* and *P. compressa* data that were generated in this study. ModelTest-NG (Darriba et al. 2019) was used to choose appropriate models of evolution for both datasets. Using the best model of evolution, maximum likelihood analyses were then performed using 1000 standard bootstraps.

VCF files from the ipyrad pipeline were further filtered using VCFtools (Danecek et al. 2011). Parameters --min-alleles 2 and --max-alleles 2 were used to filter for only biallelic loci and --mac 3 was used to remove minor allele counts < 3 as suggested by Linck and Battey (2019). The populations program of Stacks (Catchen et al. 2013) was then used to generate F-statistics. Between sites, loci were required to be in ≥6/8 sites in order to be processed. Within sites, ≥2/3 of the individuals were required to possess a locus in order for it to be processed. Additionally, maximum observed heterozygosity was restricted to ≤0.5 and an F_ST_ correction was applied such that if an F_ST_ value was not significantly different than 0, its value was set to 0. The same parameters were used to generate files for STRUCTURE (Pritchard et al. 2000) with the addition of --write_random_snp to randomly select one single nucleotide polymorphism (SNP) per locus to prevent the inclusion of linked loci. In STRUCTURE, five runs of 50,000 iterations across K=1, 2, 3, 4, 5, and 6 were performed for each species with 10,000 iterations disposed as burn-in. The analysis methods used in this study have been utilized with great success in studies of scleractinians as well as other taxa with similar issues of reticulate evolution such as American live oaks (Cavender-Bares et al. 2015).

To identify clones represented in the dataset, the script vcf_clone_detect.py (https://github.com/pimbongaerts/radseq/blob/master/vcf_clone_detect.py) was utilized to calculate pairwise genetic similarity between sampled colonies. A threshold of 95% similarity was supplied to classify samples as clonal.

### Mantel Tests

Mantel tests were performed to test for correlations between genetic distance (F_ST_) and geographic distance data for each species using ade4 (Dray and Dufour 2007). Genetic distance matrices were generated using Stacks populations and geographic distance matrices were generated from GPS coordinates using Geographic Distance Matrix Generator v. 1.2.3 (Ersts n.d.). Average temperature and salinity variability (range) was calculated at daily, weekly, monthly, and yearly time scales using 12 months of ROMS model output data (May 2018-May 2019) from the Pacific Islands Ocean Observing System (PacIOOS, pacioos.hawaii.edu). Average sea surface height was also calculated at the different time scales. Model data was downloaded for each site GPS coordinate and the respective depth that the coral samples were collected at. Additional mantel tests were also performed using data from past publications including water residence time (Lowe et al. 2009), and average pCO_2_ (Fagan and Mackenzie 2007).

## Results

### Assembly and population summary statistics

Approximately 260M reads were generated for all samples combined. Two outliers were obtained, ∼11M reads for one *M. capitata* (sample ID M7W_A) and ∼43K reads for one *P. compressa* colony (P2_C). P2_C was removed from subsequent analysis due to poor read quantity and sequence quality. For *M. capitata* and *P. compressa*, the average number of reads passing default S2 ipyrad filters was 3.02M and 2.47M, respectively. Reads were assigned to ∼109,000 high depth clusters with an average depth of 5.43 for *M. capitata* and ∼79,000 high depth clusters with an average depth of 7.94 for *P. compressa.* The final step of default filters in ipyrad resulted in 77,792 retained loci in the *M. capitata* assembly and 42,166 retained loci in the *P. compressa* assembly.

Pairwise F_ST_ values (**Table 1**) for sites 1-8 in *M. capitata* are relatively consistent between all sites, ranging from F_ST_ = 0.0452 – 0.0614, with a slight increase as one moves in a northwest-southeast direction. Pairwise F_ST_ values for sites 1-8 in *P. compressa* vary more between sites than in *M. capitata*, F_ST_ = 0.0501 – 0.1209, and in a northeast-southwest direction. In *M. capitata*, observed heterozygosity is similar for all sites while some variability is observed in *P. compressa* (**Table 2**). Inbreeding coefficient values, F_IS_, vary slightly between sites, with values for *M. capitata* ranging from 0.033-0.053 in variant sites and values for *P. compressa* ranging from 0.015-0.088.

**Table 1:** Pairwise Fst values for each sampled site. *Montipora capitata* is shown in the top diagonal and *Porites compressa* is shown in the lower diagonal.

|  | Site 1 | Site 2 | Site 3 | Site 4 | Site 5 | Site 6 | Site 7 | Site 8 |
| --- | --- | --- | --- | --- | --- | --- | --- | --- |
| Site 1 |  | 0.056 | 0.0539 | 0.0559 | 0.0556 | 0.0554 | 0.0605 | 0.0614 |
| Site 2 | 0.1034 |  | 0.0491 | 0.0521 | 0.0499 | 0.0513 | 0.0548 | 0.0573 |
| Site 3 | 0.0501 | 0.1051 |  | 0.0504 | 0.0486 | 0.0496 | 0.0548 | 0.0544 |
| Site 4 | 0.0607 | 0.1209 | 0.0511 |  | 0.0477 | 0.0474 | 0.0452 | 0.0484 |
| Site 5 | 0.0501 | 0.1009 | 0.0505 | 0.0625 |  | 0.0476 | 0.05 | 0.0515 |
| Site 6 | 0.0653 | 0.1095 | 0.0659 | 0.0781 | 0.0676 |  | 0.0483 | 0.0505 |
| Site 7 | 0.054 | 0.1175 | 0.0602 | 0.067 | 0.0596 | 0.0708 |  | 0.0488 |
| Site 8 | 0.0697 | 0.0957 | 0.0713 | 0.0826 | 0.071 | 0.0729 | 0.0732 |  |

**Table 2:** Population summary statistics for *Montipora capitata* and *Porites compressa* at sites in Kaneohe Bay. Statistics calculated with STACKS Populations v2.4. FIS=inbreeding coefficient, Obs. Hom.=Observed homozygosity, Obs. Het.=Observed heterozygosity, Exp. Hom.=Expected homozygosity, Exp. Het.=Expected heterozygosity.

|  | All Positions (Variant and Fixed) |  |  |  |  |  |  |  |  |  |  |  |  |  |  |  |
| --- | --- | --- | --- | --- | --- | --- | --- | --- | --- | --- | --- | --- | --- | --- | --- | --- |
|  | *Montipora capitata* |  |  |  |  |  |  |  | *Porites compressa* |  |  |  |  |  |  |  |
|  | Site 1 | Site 2 | Site 3 | Site 4 | Site 5 | Site 6 | Site 7 | Site 8 | Site 1 | Site 2 | Site 3 | Site 4 | Site 5 | Site 6 | Site 7 | Site 8 |
| Total Sites | 64560 | 67795 | 67509 | 65253 | 68216 | 60254 | 69121 | 68349 | 64218 | 65973 | 63876 | 49385 | 65690 | 59174 | 63471 | 62861 |
| Variant Sites | 63707 | 66864 | 66599 | 64392 | 67289 | 59489 | 68171 | 67406 | 63457 | 65211 | 63138 | 48865 | 64911 | 58510 | 62762 | 62177 |
| Private Alleles | 143 | 136 | 128 | 92 | 117 | 108 | 161 | 139 | 166 | 1529 | 322 | 107 | 203 | 309 | 231 | 466 |
| % Polymorphic Loci | 65.099 | 68.57 | 68.803 | 69.269 | 69.173 | 68.704 | 70.588 | 68.831 | 65.348 | 52.315 | 65.471 | 57.665 | 65.573 | 58.742 | 63.257 | 63.001 |
| Fis | 0.032 | 0.04 | 0.049 | 0.053 | 0.041 | 0.04 | 0.044 | 0.045 | 0.078 | 0.015 | 0.074 | 0.058 | 0.082 | 0.073 | 0.08 | 0.087 |
| Nucleotide Div. (pi) | 0.215 | 0.223 | 0.224 | 0.225 | 0.224 | 0.223 | 0.231 | 0.226 | 0.215 | 0.197 | 0.214 | 0.202 | 0.215 | 0.208 | 0.21 | 0.219 |
|  | Variant Positions |  |  |  |  |  |  |  |  |  |  |  |  |  |  |  |
|  | *Montipora capitata* |  |  |  |  |  |  |  | *Porites compressa* |  |  |  |  |  |  |  |
|  | Site 1 | Site 2 | Site 3 | Site 4 | Site 5 | Site 6 | Site 7 | Site 8 | Site 1 | Site 2 | Site 3 | Site 4 | Site 5 | Site 6 | Site 7 | Site 8 |
| Obs. Hom. | 0.796 | 0.792 | 0.795 | 0.796 | 0.792 | 0.792 | 0.786 | 0.791 | 0.816 | 0.801 | 0.815 | 0.824 | 0.817 | 0.825 | 0.822 | 0.819 |
| Standard Error | 0.001 | 0.001 | 0.001 | 0.001 | 0.001 | 0.001 | 0.001 | 0.001 | 0.001 | 0.001 | 0.001 | 0.001 | 0.001 | 0.001 | 0.001 | 0.001 |
| Obs. Het. | 0.204 | 0.208 | 0.205 | 0.204 | 0.208 | 0.208 | 0.214 | 0.209 | 0.184 | 0.199 | 0.185 | 0.176 | 0.183 | 0.175 | 0.178 | 0.181 |
| Standard Error | 0.001 | 0.001 | 0.001 | 0.001 | 0.001 | 0.001 | 0.001 | 0.001 | 0.001 | 0.001 | 0.001 | 0.001 | 0.001 | 0.001 | 0.001 | 0.001 |
| Exp. Hom. | 0.801 | 0.794 | 0.793 | 0.792 | 0.792 | 0.795 | 0.787 | 0.791 | 0.801 | 0.821 | 0.803 | 0.815 | 0.801 | 0.808 | 0.806 | 0.798 |
| Exp. Het. | 0.199 | 0.206 | 0.207 | 0.208 | 0.208 | 0.205 | 0.213 | 0.209 | 0.199 | 0.179 | 0.197 | 0.185 | 0.199 | 0.192 | 0.194 | 0.202 |
| Fis | 0.033 | 0.041 | 0.05 | 0.053 | 0.042 | 0.04 | 0.044 | 0.046 | 0.079 | 0.015 | 0.075 | 0.059 | 0.083 | 0.074 | 0.081 | 0.088 |
| Nucleotide Div. (pi) | 0.218 | 0.226 | 0.227 | 0.228 | 0.227 | 0.225 | 0.234 | 0.229 | 0.218 | 0.2 | 0.217 | 0.204 | 0.218 | 0.211 | 0.213 | 0.222 |

Mantel tests revealed significant correlation between *P. compressa* F_ST_ values and yearly average sea surface height. *M. capitata* values were significantly correlated with geographic distance, water residence time, yearly temperature range, monthly temperature range, weekly temperature range, monthly salinity range, weekly salinity range, and daily salinity range at an alpha level of p<0.05 (**Fig. 2**).

**Figure 2:**
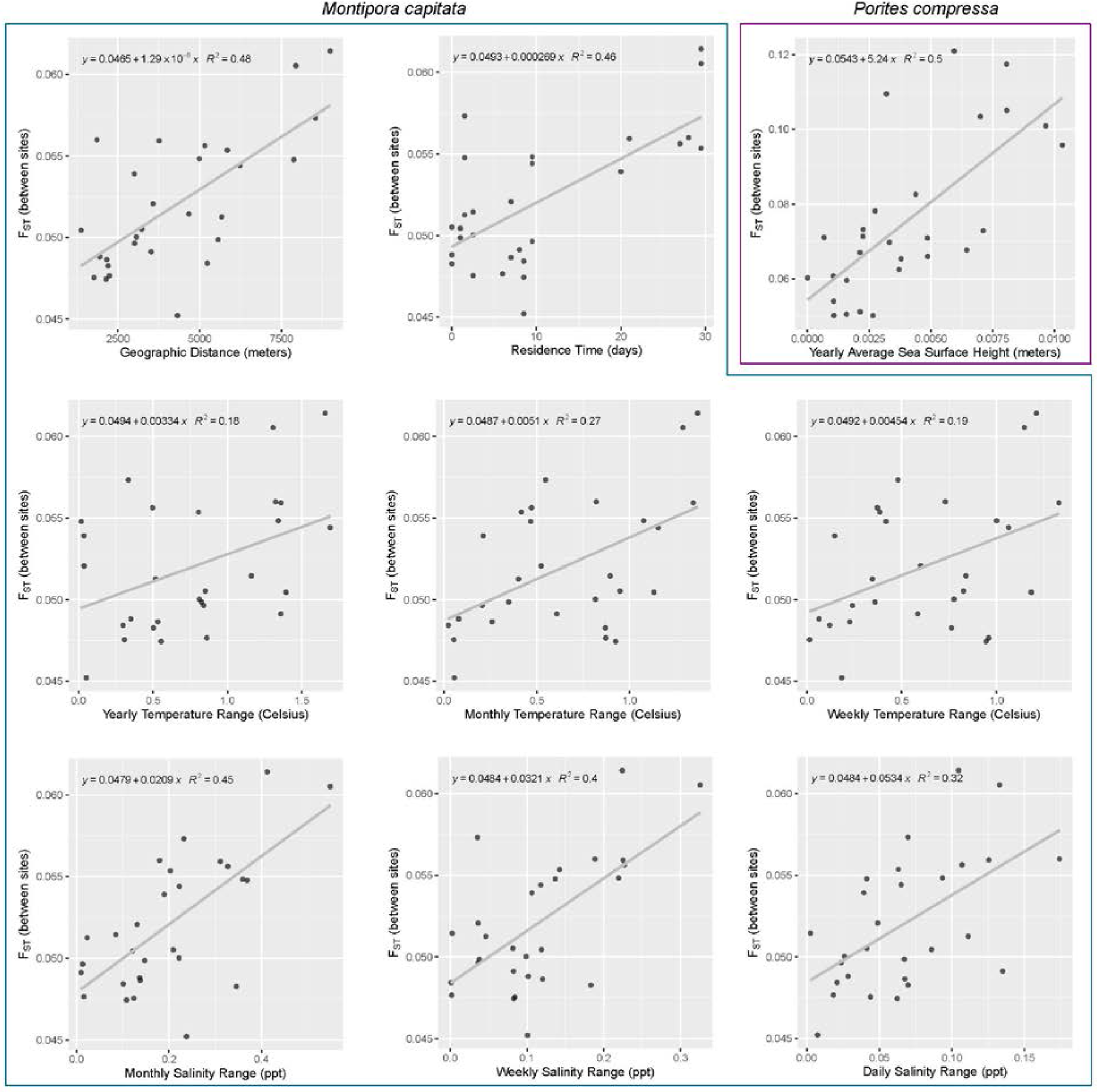
Significant Mantel tests for *Montipora capitata* and *Porites compressa* in Kaneohe Bay, Oahu. Variables shown are geographic distance (p=0.0014), residence time (p=0.0392), yearly temperature range (p=0.0223), monthly temperature range (p=0.0047), weekly temperature range (p=0.0138), monthly salinity range (p=0.0057, weekly salinity range (p=0.0128), daily salinity range (p=0.0151), and yearly average sea surface height (p=0.0267).

### Clustering Analyses

Clustering analyses were performed using STRUCTURE 2.3.4 (Pritchard et al. 2000) with the admixture model. The Evanno method (Evanno et al. 2005) as implemented in STRUCTURE Harvester (Earl and vonHoldt 2012) indicated the optimal value of K for *Montipora* samples to be K=2 while the optimal value of K for *Porites* samples was found to be K=3 (**Supplemental Figure 1**). Site-level and individual-level probability of membership for each species are shown in **Fig. 3**. The STRUCTURE analyses reveal clear population structure patterns in *Montipora* but no apparent clustering patterns amongst sampled *Porites. Montipora* samples somewhat resemble an isolation by distance scenario in which the north and south are distant geographically or environmentally from one another.

**Figure 3:**
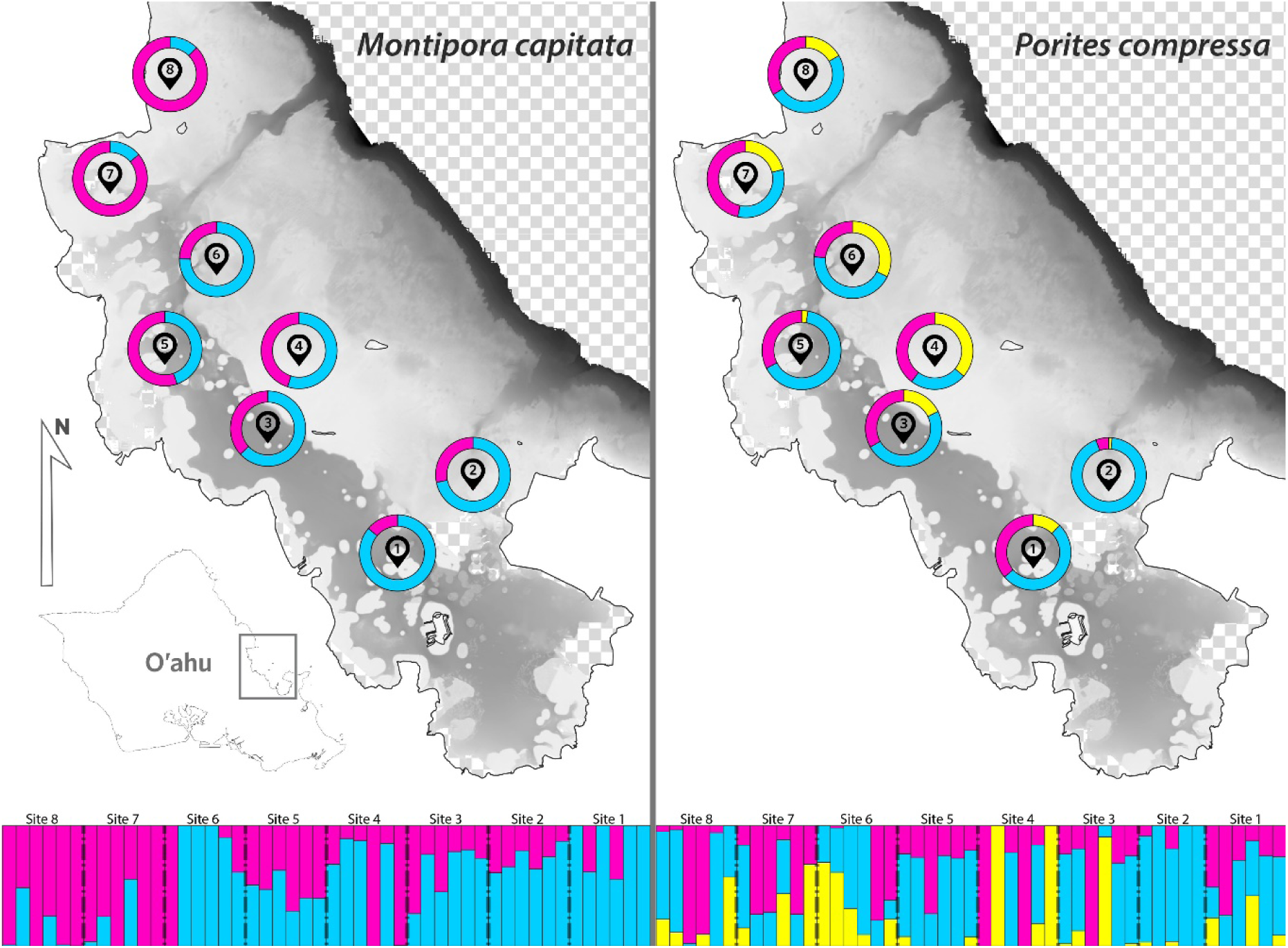
Site-level probability of membership (donut plots on map) and individual-level probability of cluster membership (bar plots at bottom) at K=2 and K=3, for *Montipora capitata* and *Porites compressa*, respectively.

### Maximum likelihood phylogeny

According to BIC, ModelTest-NG determined the best-fit model of evolution for both *M. capitata* and *P. compressa* was TVM+ASC. RAxML-NG maximum likelihood analyses of *Montipora* samples did not converge after 1000 bootstrap iterations. Analysis of *Porites* data converged after 500 bootstrap iterations. Transfer bootstrap expectation (TBE) support values were mapped to the maximum likelihood tree topology and phylogenies for both species are reported in **Fig. 4**. The unconverged *M. capitata* tree was poorly supported and had strong support only at tips. The *P. compressa* tree was strongly supported at both basal and terminal nodes.

**Figure 4:**
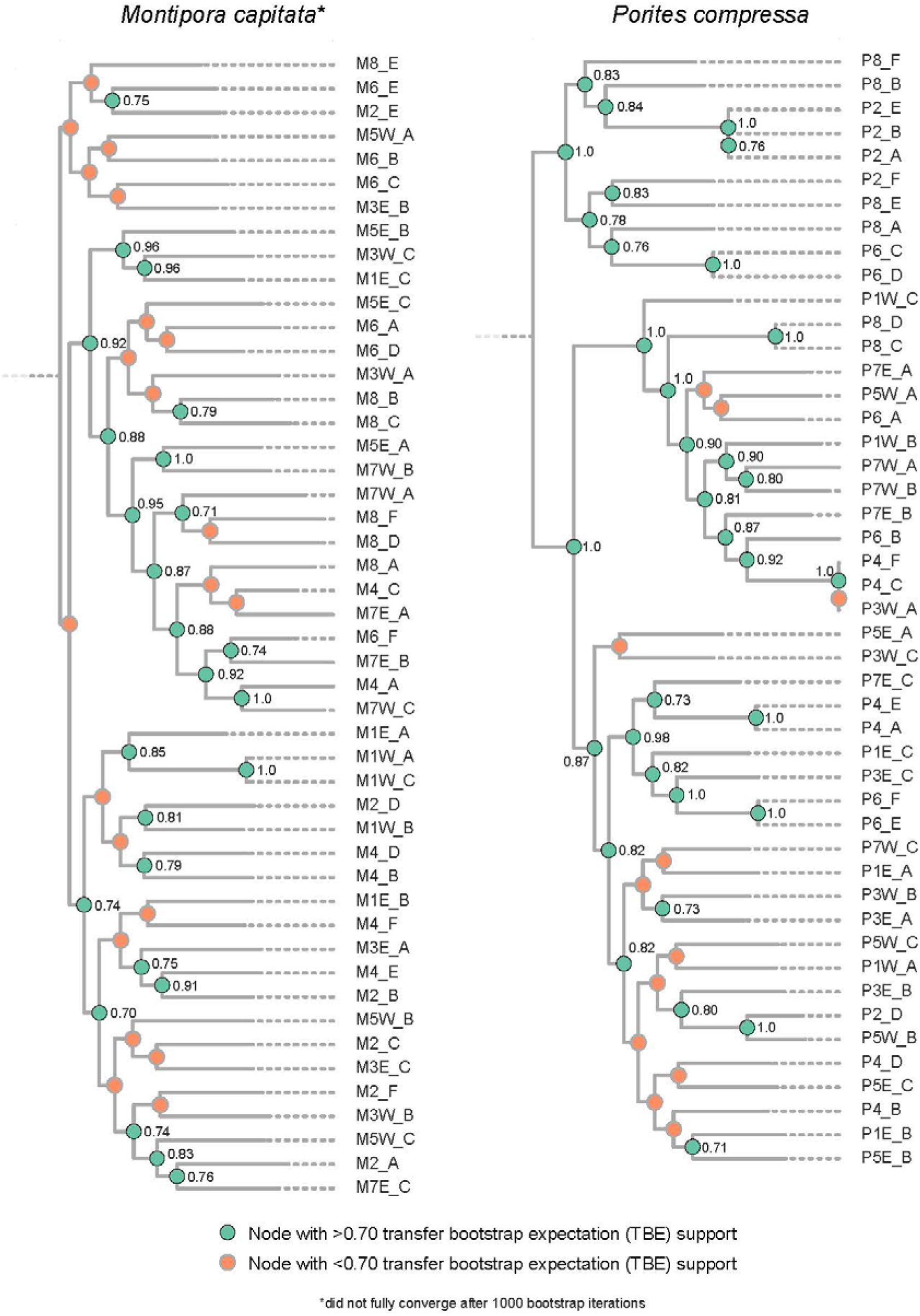
Population phylogenies of *Montipora capitata* (left) and *Porites compressa* (right). *M. capitata* phylogeny did not reach convergence after 1000 standard bootstrap iterations in RAxML-NG. *M. capitata* tree is rooted by *M. spumosa* and *P. compressa* tree is rooted by *P. lutea*.

### Clonal groups

Analysis of pairwise percent similarity between individuals showed an average genetic similarity of 78.64% with a standard deviation (SD) of 1.05% in *Montipora capitata* and 77.37% with a SD of 2.95% in *Porites compressa*. Distribution of values was unimodal in *M. capitata* and bimodal in *P. compressa*. Clonal groups were identified by a threshold of 95%, following the logic that clonal individuals should be nearly 100% identical. In *M. capitata*, this present study found only one clonal pair of colonies, existing at site 1, adjacent to Coconut Island. In *P. compressa*, two clonal triplets and four clonal pairs were detected (**Fig. 5**). Spatially, these clonal groupings occurred predominantly at outer bay sites 2, 4, 6, and 8, with only one inner bay colony, P3W_A, being represented as part of a clonal group. Clonal colonies made up the majority of samples recovered in sites 2, 4, and 6. At these three sites, a total of 17 genotypes were expected but only 11 were detected using our sampling design and ddRAD methods.

**Figure 5:**
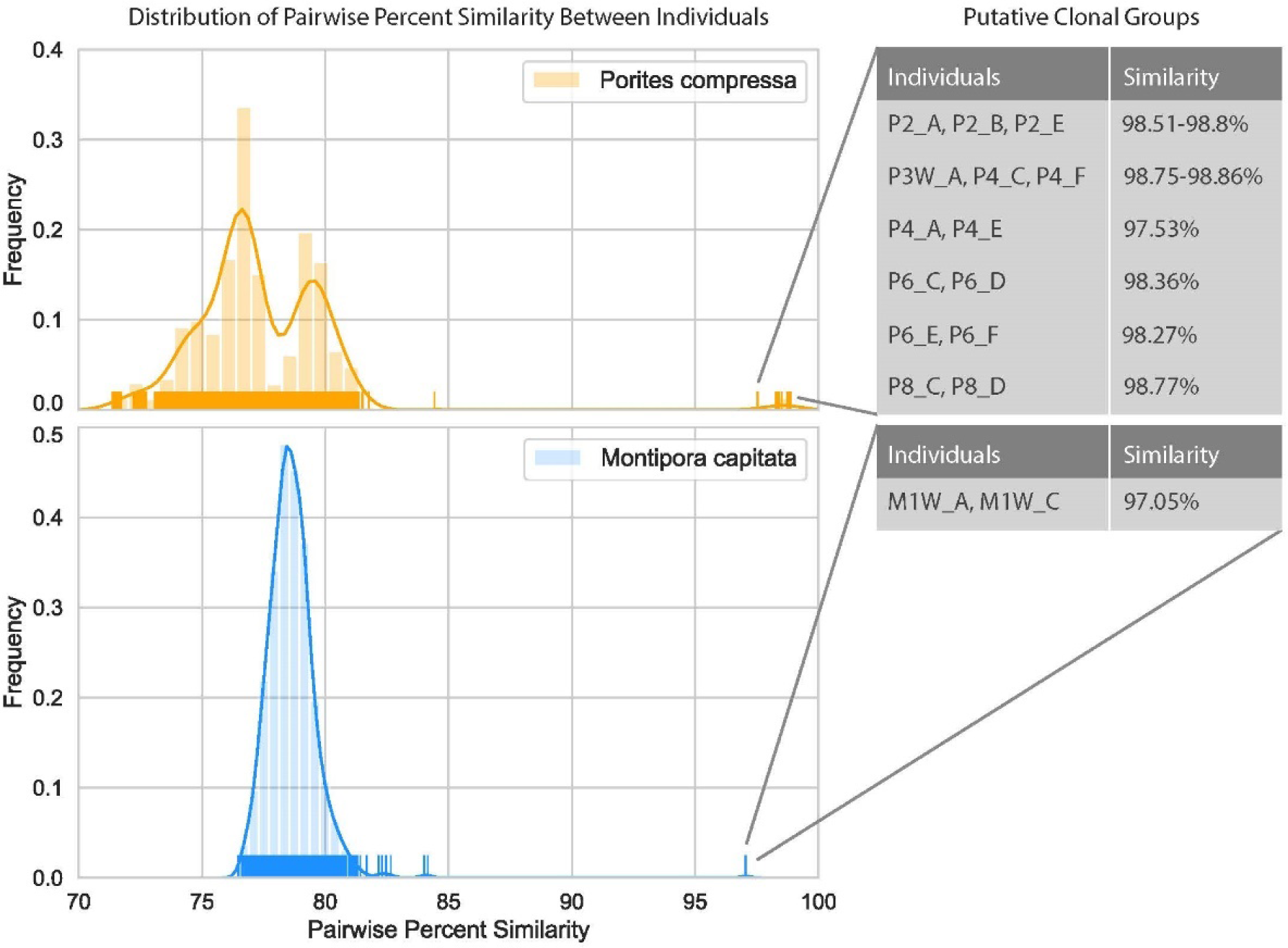
Distribution of pairwise percent similarity values for *Montipora capitata* and *Porites compressa*. Putative clonal groups (as suggested by distributions and a 95% threshold) are shown on the right.

## Discussion

### Patterns of population structure

This study found signals of population structure in *M. capitata* on a very fine-scale seascape. Such a finding is unusual for both the system – broadcast spawning marine organisms – and the spatial scale. Despite findings, this study cannot discern what drives spatial patterns of structure. In this study, mantel tests revealed significant correlations between *M. capitata* F_ST_ values and geographic distance, water residence time, and temperature and salinity variability at various temporal scales. Because all of these variables are linked, it is difficult to discern which variable or multiple variables drive the patterns of structure. However, global and local analyses have found high-frequency temperature variability to be the most influential factor in predicting bleaching occurrence and percent coral coverage (Soto et al. 2011, Carilli et al. 2012, Safaie et al. 2018). This present study, combined with results of past studies, suggest that temperature variability may be playing a role in population structure of *M. capitata* in KB. However, it is worth noting that these studies focused on either a) all reef regions globally or b) forereef systems locally and may not have captured the effect that salinity can have on fine-scale lagoonal systems such as KB.

In addition to parameters such as temperature and salinity, physical barriers such as ocean currents may partially explain patterns of structure in corals. The presence of the Mokapu peninsula at the eastern side of KB causes ocean currents to split the bay into a northern and southern section during the course of the coral spawning period (Richmond and Hunter 1990, Padilla-Gamiño and Gates 2012) (**Fig. 6**). Because water cannot easily escape the sheltered southern portion of the bay, the north and south are distinct in their residence times. In the north, water remains in the bay for ≤5 days while water in the south can remain in the bay for ≥15 days (Lowe et al. 2009). The distinct zones of residence in KB may partially drive patterns of settlement and population structure that we observe in this study. Acroporids have short times to settlement, typically ranging between 1-6 days (Jones et al. 2015). Due to residence times ≥15 days, southern bay sites would be restricted primarily to self-recruitment of acroporid larvae. Sites in the north experience shorter water residence times than typical time-to-settlement durations of acroporids and, thus, can export and exchange larvae with peripheral habitats. It is worth noting that the models predicting residence time in Lowe et. al were not specific to the coral spawning period.

**Figure 6:**
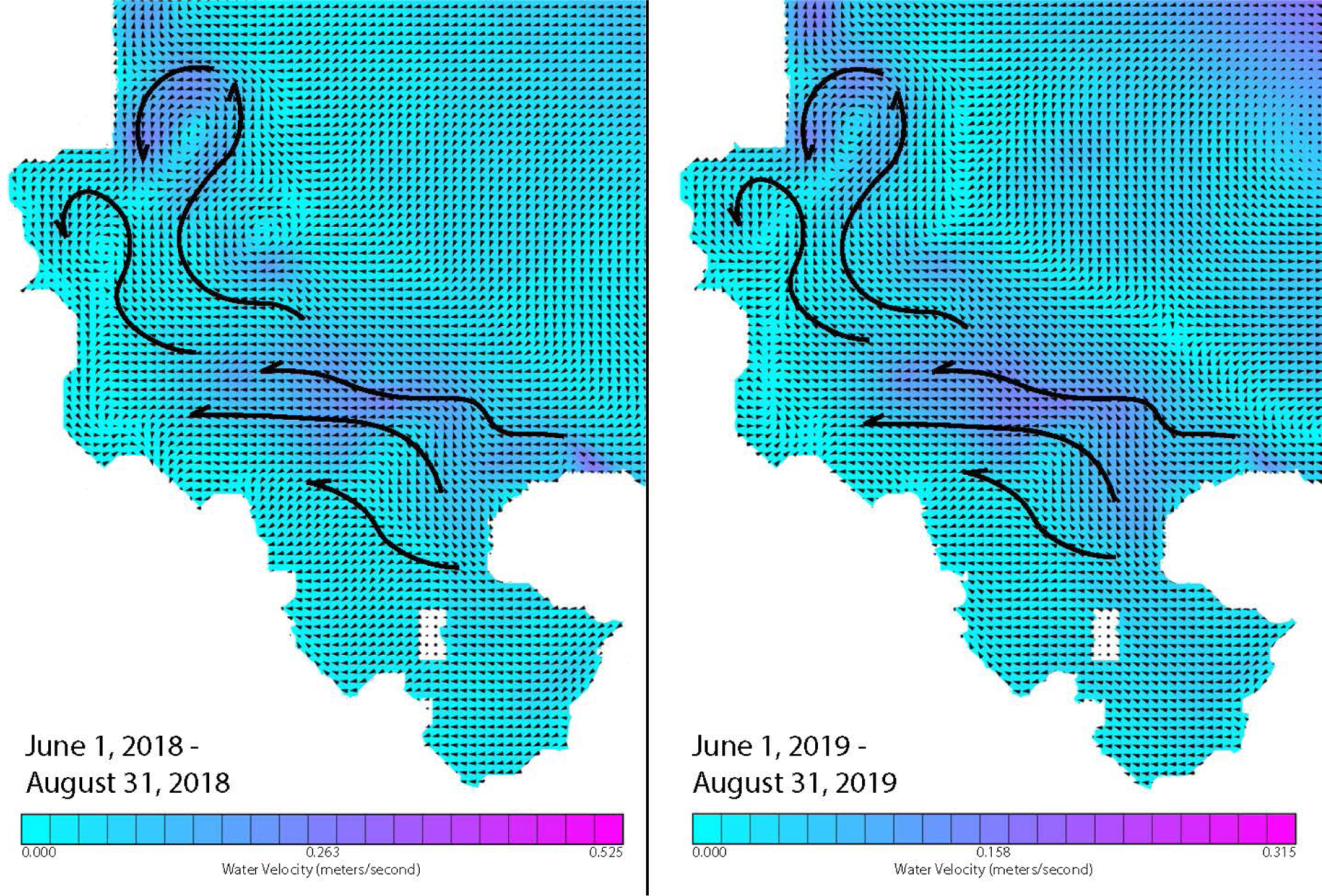
A map of surface currents in Kaneohe Bay during the coral spawning period (June-August) of Hawaii for 2018 and 2019. Data sourced from the Pacific Islands Ocean Observing System ROMS model.

No apparent spatial patterns of population structure that align with temperature, salinity, or residence time zones were detected in *P. compressa*. Carilli et al. found historical temperature variability to be an important predictor of bleaching and partial mortalities in massive *Porites* spp. (2012). However, Mantel tests utilized this present study did not suggest temperature to be an important factor. Residence time zones may not be important as typical pelagic larval duration may exceed the longest residence times found in KB. Studies of *Porites* larval duration and reproductive success are rare, but other taxa with similar massive morphologies show drastically longer larval longevities than those of acroporids (Graham et al. 2008). *P. compressa* has preference for sheltered lagoons and has been shown in models and surveys to not hold up to significant wave action (Rodgers et al. 2004, Franklin et al. 2013). Correlations between *P. compressa* F_ST_ values and yearly average sea surface height found in this present study align with these models and surveys.

It is worth noting that in broader phylogenetic studies, *P. compressa* and *P. lobata* do not form distinct clades and morphologically identified *P. compressa* may fall in *P. lobata* dominated clades, and vice versa (Forsman et al. 2017). It is possible that cryptic species may be obfuscating patterns of structure and F_ST_-environment correlations. A majority of *P. compressa* in outer bay sites 2, 6, and 8 form a strongly supported clade at the base of the phylogeny. Sites 2 and 6 represent regions of overlap for modeled coral range and abundance of morphologically-identified *P. compressa* and *P. lobata* (Franklin et al. 2013). It is plausible that this strongly-supported basal clade is present due to cryptic species or hybridization and introgression between species. Additional evidence of cryptic species or reticulate evolution can be found in distributions of percent pairwise similarity between *P. compressa* individuals (**Fig. 5**). Bimodal distributions of percent pairwise similarity may suggest populations of a single species undergoing disruptive selection or two separate taxa being represented in the genetic dataset.

### Spatial distribution of clonality

Past work to quantify prevalence of clonality in *P. compressa* found that regions with histories of disturbance contained proportionally fewer clonal colonies compared to those of sexual origin (Hunter 1993). Specifically, less disturbed locations were more likely to be space-limited and recruits of sexual origin would struggle to settle. In disturbed locations, openings would commonly exist on the benthic substrate and allow for recruitment of larvae. Although the methodology of our study was not designed specifically to address the question of clonality, we show that clonality is much more prevalent in locations in the outer bay. These regions experience high energy swells and are prone to storm surge which can fragment corals or dislodge natural subspheroidal coralliths (Glynn 1974, Roff 2008, Capel et al. 2012). However, this phenomenon appears to be biased toward *P. compressa* as few clones were detected in *M. capitata*. It is possible that clonal colony formation may be more prevalent in *P. compressa* due to fundamental differences in life history traits. When natural growth rate is slow, as in *Porites* spp., new colonies may be given a “jump start” by growing from wave-induced fragments or rolling coralliths, rather than having to grow from larvae. Additionally, hydrodynamic studies have predicted that nudibranch larvae can settle only on sheltered areas of reefs because wave action can dislodge settling larvae (Reidenbach et al. 2009). Perhaps this same mechanism is at work in *P. compressa* and is what drives fragmentary reproduction to be favored over sexually produced larvae in reefs with high wave action. In *Montipora* spp., growth is fast and fragmentation may not offer significant benefits over reproduction that occurs sexually. Despite the advantages of clonal colony formation, asexual reproduction lowers per-population genetic diversity. If storm frequency and intensity are to increase as suggested by climate models of Hawaii (Murakami et al. 2013), it is possible that population genetic diversity of *P. compressa* populations will decrease, regardless of other pressures such as temperature increases and sedimentation.

### Phylogeographic and population structure patterns in relation to bleaching extent and recovery

Although we cannot necessarily tease apart the causality of genetic patterns in this study, it is worth noting parallels between our results and past bleaching events in KB. A study of the 1996 bleaching event focused on sites with >90% coral cover and these sites contained >90% *P. compressa* by percent cover (Jokiel and Brown 2004). As such, we cannot compare this study to our findings of *M. capitata*. During this 1996 bleaching event, surveys were performed immediately adjacent to our sites 1, 2, 3, 5, and 6. In this set of surveys, it was found that sites in the inner bay (adjacent to sites 1, 3, and 5) encountered extensive bleaching while outer bay sites (adjacent to sites 2 and 6) remained mostly unscathed. Our data show that four out of five individuals at site 2 and two out of six individuals at site 6 are members of a strongly supported basal clade in the *P. compressa* phylogeny. While there are other factors that are likely to drive bleaching response, we show in this present study that there is also some level of genetic divergence between populations that exhibited different responses to bleaching thresholds.

In the bleaching event of 2014, the symbiont community composition of *M. capitata* colonies was monitored as bleaching progressed as well as during recovery after the event (Cunning et al. 2016). This study only included colonies in the inner bay, adjacent to our sites 1, 3, and 5. Cunning et al. (2016) found that bleaching response in *M. capitata* was significantly associated with dominant symbiont clade but that the symbiont communities did not cluster spatially. Additionally, it was found that recovery rates increased the further north individuals were within the bay. Our study shows that there is population structure along a north-south gradient within KB and that this aligns with the spatial distribution of post-bleaching recovery rates.

It is important to note that these bleaching events were fundamentally different, as discussed by Bahr et al. (2017). The timing and environmental conditions both played a key role in their extent, severity, and mortality rates. Despite the spatial and temporal differences between events, we believe that past studies, combined with the genetic results of this study, provide some support that the population genetics of the coral host likely acts synergistically with environmental variables, stochastic events, and symbiont community compositions to produce a bleaching response.

## Acknowledgements

Thank you to the brightest and most resilient field assistant, Montana Airey. No hurricane can stop us. Thank you to the Drew Lab for the support and help on my thesis research. I acknowledge the State of Hawaii for permitting this work and the E3B department at Columbia University, the Society of Systematic Biologists Graduate Student Research Award, and the Earth Institute Travel Grant for providing funding for this research.

## Data Accessibility

Raw sequence reads generated in this study are deposited under NCBI BioProject accession number PRJNA544861. Jupyter notebooks used to process and analyze data are publicly accessible in the following GitHub repository: https://github.com/mistergroot/kbaygen

## Supplemental materials

**Supplemental Table 1:**
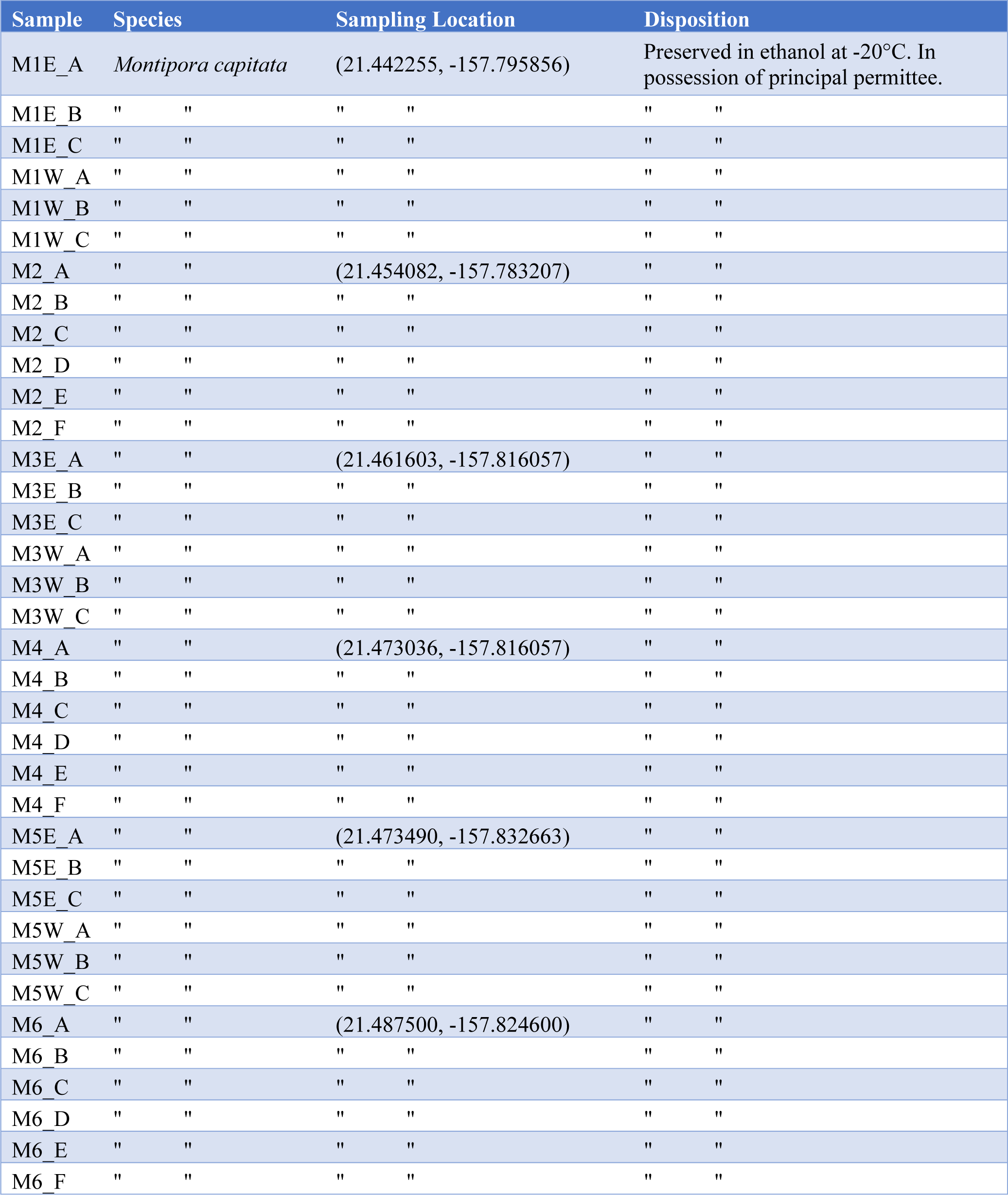
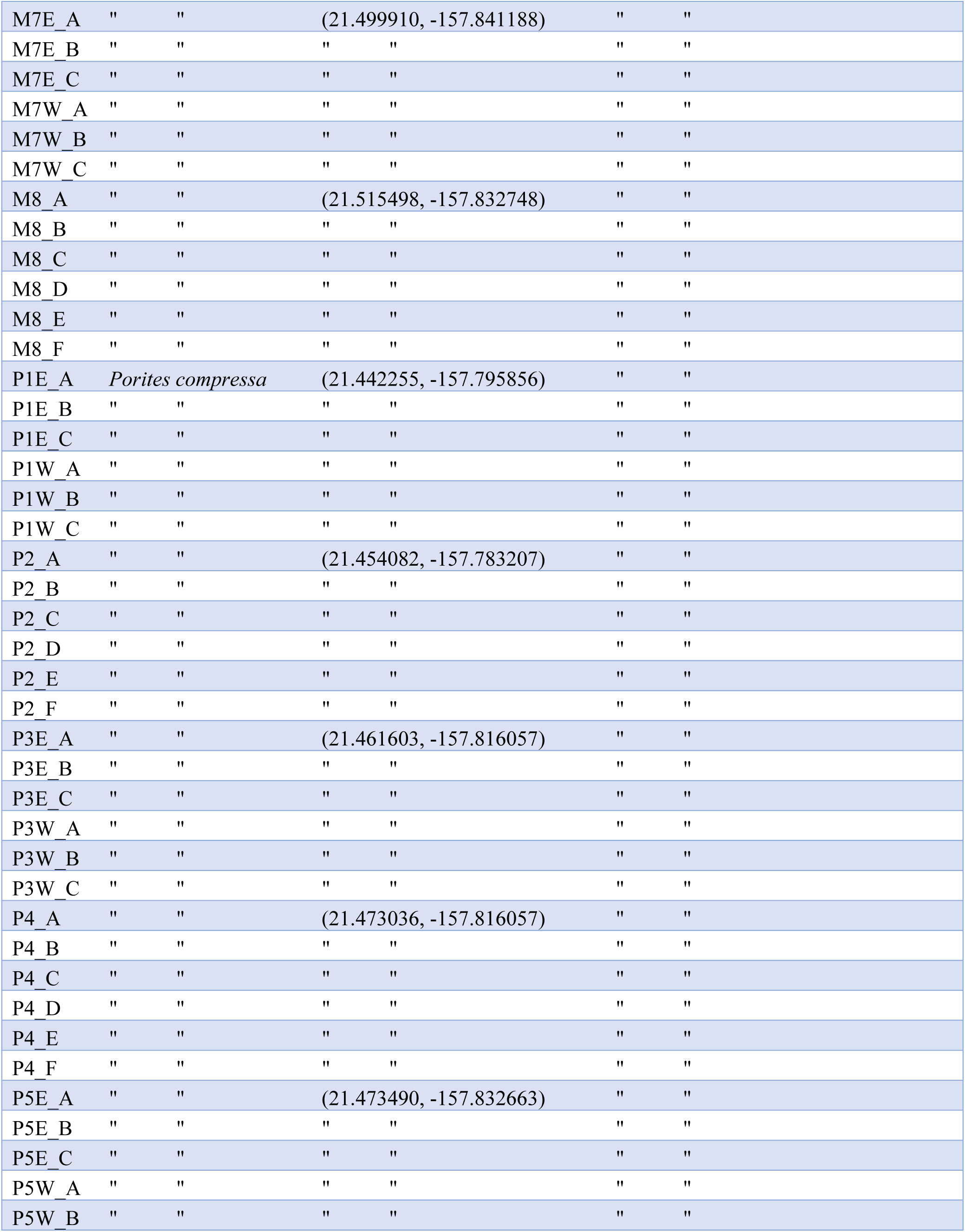
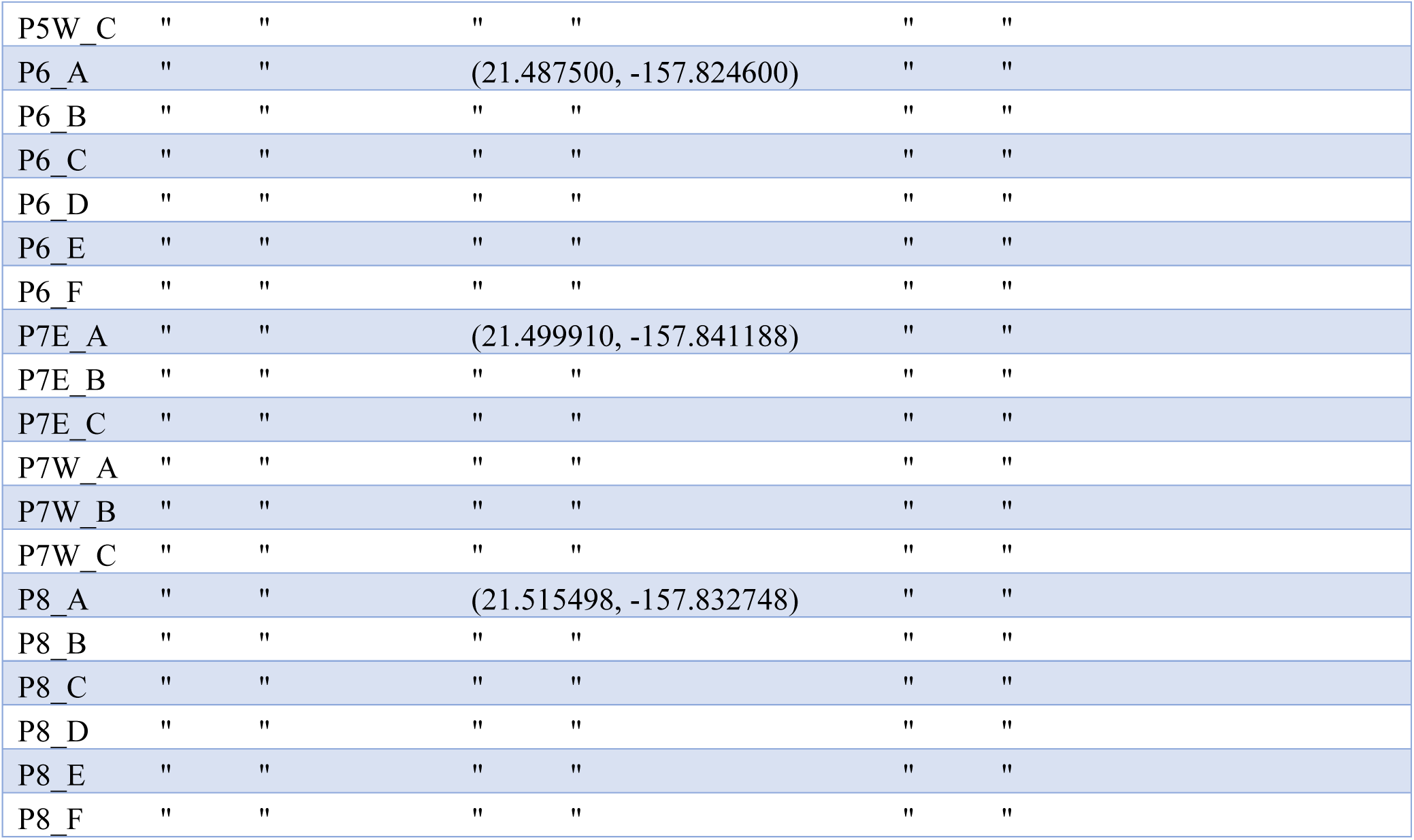
Sample inventory (as of 12/10/2019) and sampling coordinates of coral colony samples obtained from Kaneohe Bay, Oahu.

**Supplemental Figure 1:**
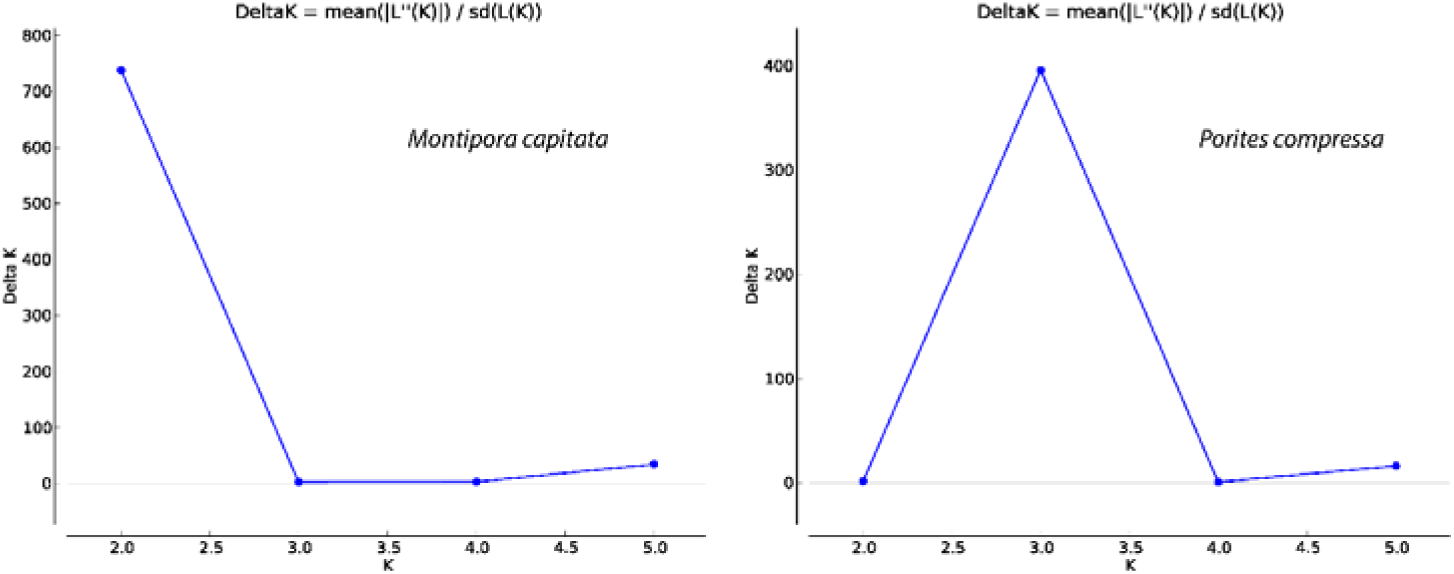
DeltaK values for *Montipora capitata* and *Porites compressa* as evaluated by StructureHarvester.

## References

Bahr, K. D., P. L. Jokiel, and K. S. Rodgers. 2015a. The 2014 coral bleaching and freshwater flood events in Kāne‘ohe Bay, Hawai‘i. PeerJ 3:e1136.

Bahr, K. D., P. L. Jokiel, and R. J. Toonen. 2015b. The unnatural history of Kāne’ohe Bay: coral reef resilience in the face of centuries of anthropogenic impacts. PeerJ 3:e950.

Bahr, K. D., K. S. Rodgers, and P. L. Jokiel. 2017. Impact of Three Bleaching Events on the Reef Resiliency of Kāne’ohe Bay, Hawai’i. Frontiers in Marine Science 4.

Bourne, D., Y. Iida, S. Uthicke, and C. Smith-Keune. 2008. Changes in coral-associated microbial communities during a bleaching event. The ISME Journal 2:350–363.

Capel, K., B. Segal, P. Bertuol, and A. Lindner. 2012. Corallith beds at the edge of the tropical South Atlantic Fig. Coral Reefs 31.

Carilli, J., S. D. Donner, and A. C. Hartmann. 2012. Historical temperature variability affects coral response to heat stress. PLoS ONE 7:1–9.

Catchen, J., P. A. Hohenlohe, S. Bassham, A. Amores, and W. A. Cresko. 2013. Stacks: an analysis tool set for population genomics. Molecular ecology 22:3124–40.

Cavender-Bares, J., A. González-Rodríguez, D. A. R. Eaton, A. A. L. Hipp, A. Beulke, and P. S. Manos. 2015. Phylogeny and biogeography of the american live oaks (Quercus subsection Virentes): A genomic and population genetics approach. Molecular Ecology 24:3668–3687.

Chafin, T. K., B. T. Martin, S. M. Mussmann, M. R. Douglas, and M. E. Douglas. 2018. FRAGMATIC: in silico locus prediction and its utility in optimizing ddRADseq projects. Conservation Genetics Resources 10:325–328.

Cheung, W. W. L., V. W. Y. Lam, J. L. Sarmiento, K. Kearney, R. Watson, and D. Pauly. 2009. Projecting global marine biodiversity impacts under climate change scenarios. Fish and Fisheries 10:235–251.

Consortium, R. 2020. 2015. The ReFuGe 2020 Consortium — using “omics” approaches to explore the adaptability and resilience of coral holobionts to environmental change. Frontiers in Marine Science 2.

Cunning, R., R. Ritson-Williams, and R. D. Gates. 2016. Patterns of bleaching and recovery of Montipora capitata in Kane’ohe Bay, Hawai’i, USA. Marine Ecology Progress Series 551:131–139.

Danecek, P., A. Auton, G. Abecasis, C. A. Albers, E. Banks, M. A. DePristo, R. E. Handsaker, G. Lunter, G. T. Marth, S. T. Sherry, G. McVean, and R. Durbin. 2011. The variant call format and VCFtools. Bioinformatics 27:2156–2158.

Darriba, D., D. Posada, T. Flouri, A. M. Kozlov, A. Stamatakis, and B. Morel. 2019. ModelTest-NG: A New and Scalable Tool for the Selection of DNA and Protein Evolutionary Models. Molecular Biology and Evolution:1–4.

Devlin-Durante, M. K., and I. B. Baums. 2017. Genome-wide survey of single-nucleotide polymorphisms reveals fine-scale population structure and signs of selection in the threatened Caribbean elkhorn coral, *Acropora palmata*. PeerJ 5:e4077.

Dray, S., and A.-B. Dufour. 2007. The ade4 package: Implementing the duality diagram for ecologists. Journal of Statistical Software 22.

Drury, C., K. E. Dale, J. M. Panlilio, S. V. Miller, D. Lirman, E. A. Larson, E. Bartels, D. L. Crawford, and M. F. Oleksiak. 2016. Genomic variation among populations of threatened coral: Acropora cervicornis. BMC Genomics 17:1–14.

Drury, C., S. Schopmeyer, E. Goergen, E. Bartels, K. Nedimyer, M. Johnson, K. Maxwell, V. Galvan, C. Manfrino, and D. Lirman. 2017. Genomic patterns in Acropora cervicornis show extensive population structure and variable genetic diversity. Ecology and Evolution 7:6188–6200.

Earl, D. A., and B. M. vonHoldt. 2012. STRUCTURE HARVESTER: A website and program for visualizing STRUCTURE output and implementing the Evanno method. Conservation Genetics Resources 4:359–361.

Eaton, D. A. R. 2014. PyRAD: Assembly of de novo RADseq loci for phylogenetic analyses. Bioinformatics 30:1844–1849.

Ersts, P. J. (n.d.). Geographic distance matrix generator (version 1.2.3). American Museum of Natural History, Center for Biodiversity and Conservation.

Evanno, G., S. Regnaut, and J. Goudet. 2005. Detecting the number of clusters of individuals using the software STRUCTURE: A simulation study. Molecular Ecology 14:2611–2620.

Fagan, K. E., and F. T. Mackenzie. 2007. Air-sea CO2 exchange in a subtropical estuarine-coral reef system, Kaneohe Bay, Oahu, Hawaii. Marine Chemistry 106:174–191.

Forsman, Z. H., I. S. S. Knapp, K. Tisthammer, D. A. R. Eaton, M. Belcaid, and R. J. Toonen. 2017. Coral hybridization or phenotypic variation? Genomic data reveal gene flow between Porites lobata and P. Compressa. Molecular Phylogenetics and Evolution 111:132–148.

Franklin, E. C., P. L. Jokiel, and M. J. Donahue. 2013. Predictive modeling of coral distribution and abundance in the Hawaiian Islands. Marine Ecology Progress Series 481:121–132.

Gladfelter, E. H., R. K. Monahan, and W. B. Gladfelter. 1978. Growth rates of five reef-building corals in the northeastern Caribbean. Bulletin of Marine Science 28:728–734.

Glynn, P. W. 1974. Rolling stones among the Scleractinia: Mobile coralliths in the Gulf of Panama. Page Second International Coral Reef Symposium.

Graham, E. M., A. H. Baird, and S. R. Connolly. 2008. Survival dynamics of scleractinian coral larvae and implications for dispersal. Coral Reefs 27:529–539.

Hughes, T. P. P., A. H. Baird, D. R. Bellwood, M. Card, S. R. Connolly, C. Folke, R. Grosberg, O. Hoegh-Guldberg, J. B. C. Jackson, J. Kleypas, J. M. Lough, P. Marshall, M. Nyström, S. R. Palumbi, J. M. Pandolfi, B. Rosen, and J. Roughgarden. 2003. Climate change, human impacts, and the resilience of coral reefs. Science 301:929–33.

Hunter, C. L. 1993. Genotypic Variation and Clonal Structure in Coral Populations with Different Disturbance Histories. Evolution 47:1213–1228.

Huston, M. 1985. Variation in coral growth rates with depth at Discovery Bay, Jamaica. Coral Reefs 4:19–25.

Jokiel, P. L., and E. K. Brown. 2004. Global warming, regional trends and inshore environmental conditions influence coral bleaching in Hawaii. Global Change Biology 10:1627–1641.

Jones, R. J., O. Hoegh-Guldberg, A. W. D. Larkum, and U. Schreiber. 1998. Temperature-induced bleaching of corals begins with impairment of the CO2 fixation mechanism in zooxanthellae. Plant, Cell and Environnment 21:1219–1230.

Jones, R., G. F. Ricardo, and A. P. Negri. 2015. Effects of sediments on the reproductive cycle of corals. Marine Pollution Bulletin 100:13–33.

Kenkel, C. D., G. Goodbody-Gringley, D. Caillaud, S. W. Davies, E. Bartels, and M. V. Matz. 2013. Evidence for a host role in thermotolerance divergence between populations of the mustard hill coral (Porites astreoides) from different reef environments. Molecular Ecology 22:4335–4348.

Kenkel, C. D., and M. V Matz. 2016. Gene expression plasticity as a mechanism of coral adaptation to a variable environment. Nature Ecology and Evolution 1.

Kozlov, A. M., D. Darriba, and A. Stamatakis. 2019. RAxML-NG: a fast, scalable and user-friendly tool for maximum likelihood phylogenetic inference. Bioinformatics:1–3.

Li, J., Q. Chen, L.-J. Long, J.-D. Dong, J. Yang, and S. Zhang. 2015. Bacterial dynamics within the mucus, tissue and skeleton of the coral Porites lutea during different seasons. Scientific Reports 4:7320.

Liew, Y. J., M. Aranda, and C. R. Voolstra. 2016. Reefgenomics.Org - a repository for marine genomics data. Database.

Linck, E., and C. J. Battey. 2019. Minor allele frequency thresholds strongly affect population structure inference with genomic datasets. Molecular Ecology Resources:0–2.

Lowe, R. J., J. L. Falter, S. G. Monismith, and M. J. Atkinson. 2009. A numerical study of circulation in a coastal reef-lagoon system. Journal of Geophysical Research 114.

Murakami, H., B. Wang, T. Li, and A. Kitoh. 2013. Projected increase in tropical cyclones near Hawaii. Nature Climate Change 3:749–754.

Oliver, J. K. 1984. Intra-colony Variation in the Growth of Acropora formosa: Extension Rates and Skeletal Structure of White (Zooxanthellae-free) and Brown-Tipped Branches. Coral Reefs 3:139–147.

Padilla-Gamiño, J. L., and R. D. Gates. 2012. Spawning dynamics in the Hawaiian reef-building coral Montipora capitata. Marine Ecology Progress Series 449:145–160.

Peterson, B. K., J. N. Weber, E. H. Kay, H. S. Fisher, and H. E. Hoekstra. 2012. Double digest RADseq: An inexpensive method for de novo SNP discovery and genotyping in model and non-model species. PLoS ONE 7.

Pritchard, J. K., M. Stephens, and P. Donnelly. 2000. Inference of population structure using multilocus genotype data. Genetics 155:945–959.

Putnam, H. M., M. Stat, X. Pochon, and R. D. Gates. 2012. Endosymbiotic flexibility associates with environmental sensitivity in scleractinian corals. Proceedings of the Royal Society B: Biological Sciences 279:4352–4361.

Reidenbach, M. A., J. R. Koseff, and M. A. R. Koehl. 2009. Hydrodynamic forces on larvae affect their settlement on coral reefs in turbulent, wavedriven flow. Limnology and Oceanography 54:318–330.

Reshef, L., O. Koren, Y. Loya, I. Zilber-Rosenberg, and E. Rosenberg. 2006. The Coral Probiotic Hypothesis. Environmental Microbiology 8:2068–2073.

Richmond, R., and C. Hunter. 1990. Reproduction and recruitment of corals: comparisons among the Caribbean, the Tropical Pacific, and the Red Sea. Marine Ecology Progress Series 60:185–203.

Rodgers, K., M. E. Field, P. L. Jokiel, E. K. Brown, and C. D. Storlazzi. 2004. A model for wave control on coral breakage and species distribution in the Hawaiian Islands. Coral Reefs 24:43–55.

Roff, G. 2008. Corals on the move: morphological and reproductive strategies of reef flat coralliths. Coral Reefs 23:343–344.

Rosenberg, E., A. Kushmaro, E. Kramarsky-Winter, E. Banin, and L. Yossi. 2009. The role of microorganisms in coral bleaching. The ISME Journal 3:139–146.

Safaie, A., N. J. Silbiger, T. R. Mcclanahan, G. Pawlak, D. J. Barshis, J. L. Hench, J. S. Rogers, G. J. Williams, and K. A. Davis. 2018. High frequency temperature variability reduces the risk of coral bleaching. Nature Communications 2018:1–12.

Shumaker, A., H. M. Putnam, H. Qiu, D. C. Price, E. Zelzion, A. Harel, N. E. Wagner, R. D. Gates, H. S. Yoon, and D. Bhattacharya. 2019. Genome analysis of the rice coral Montipora capitata. Scientific Reports 9:2571.

Soto, I. M., F. E. Muller Karger, P. Hallock, and C. Hu. 2011. Sea Surface Temperature Variability in the Florida Keys and Its Relationship to Coral Cover. Journal of Marine Biology 2011:1–10.

Stat, M., C. E. Bird, X. Pochon, L. Chasqui, L. J. Chauka, G. T. Concepcion, D. Logan, M. Takabayashi, R. J. Toonen, and R. D. Gates. 2011. Variation in Symbiodinium ITS2 sequence assemblages among coral colonies. PLoS ONE 6.

Stat, M., X. Pochon, E. C. Franklin, J. F. Bruno, K. S. Casey, E. R. Selig, and R. D. Gates. 2013. The distribution of the thermally tolerant symbiont lineage (Symbiodinium clade D) in corals from Hawaii: Correlations with host and the history of ocean thermal stress. Ecology and Evolution 3:1317–1329.

Warner, M. E., W. K. Fitt, and G. W. Schmidt. 1999. Damage to photosystem II in symbiotic dinoflagellates: A determinant of coral bleaching. Proceedings of the National Academy of Sciences 96:8007–8012.

